# Molecular regulation of retinal regeneration is context specific

**DOI:** 10.1101/2023.11.20.567904

**Authors:** Kevin Emmerich, John Hageter, Thanh Hoang, Pin Lyu, Abigail V. Sharrock, Anneliese Ceisel, James Thierer, Zeeshaan Chunawala, Saumya Nimmagadda, Isabella Palazzo, Frazer Matthews, Liyun Zhang, David T. White, Catalina Rodriquez, Gianna Graziano, Patrick Marcos, Adam May, Tim Mulligan, Barak Reibman, Meera T. Saxena, David F. Ackerley, Jiang Qian, Seth Blackshaw, Eric Horstick, Jeff S. Mumm

## Abstract

Many genes are known to regulate retinal regeneration following widespread tissue damage. Conversely, genes controlling regeneration following limited retinal cell loss, akin to disease conditions, are undefined. Combining a novel retinal ganglion cell (RGC) ablation-based glaucoma model, single cell omics, and rapid CRISPR/Cas9-based knockout methods to screen 100 genes, we identified 18 effectors of RGC regeneration kinetics. Surprisingly, 32 of 33 previously known/implicated regulators of retinal tissue regeneration were not required for RGC replacement; 7 knockouts accelerated regeneration, including *sox2, olig2,* and *ascl1a*. Mechanistic analyses revealed loss of *ascl1a* increased “fate bias”, the propensity of progenitors to produce RGCs. These data demonstrate plasticity and context-specificity in how genes function to control regeneration, insights that could help to advance disease-tailored therapeutics for replacing lost retinal cells.

**One sentence summary:** We discovered eighteen genes that regulate the regeneration of retinal ganglion cells in zebrafish.

## Introduction

Insights on genes that regulate retinal regeneration have come almost exclusively from widespread damage paradigms(1–4). In contrast, factors controlling regeneration in the context of selective retinal cell loss, a hallmark of disease, are largely unknown. Upon retinal damage/cell loss, zebrafish Müller glia (MG) cells dedifferentiate to a stem-like state and divide asymmetrically to produce a Muller glia-derived progenitor cell (MGPC). MGPCs then proliferate before differentiating to replace lost cells. A recent comparative study of two widespread damage paradigms in zebrafish revealed this process is exquisitely tuned to the nature of the retinal injury, showing that MGPC proliferation rates and cell fate decisions were directly correlated to the numbers and types of cell lost(4). Similarly, prior studies of selective retinal cell ablation paradigms had shown that MGPCs exhibit fate bias, i.e., preferentially giving rise to the lost retinal cell type(4–7). These findings raise the possibility that mechanisms governing retinal regenerative processes are context specific; however this has yet to be rigorously tested.

To explore mechanisms regulating regeneration in the context of selective retinal cell loss, we used single cell transcriptomics and large-scale genetic screening to identify genes altering RGC regeneration kinetics following RGC ablation. We found: (1) largely unique transcriptomic signatures of reactive MG signatures following RGC ablation versus widespread damage paradigms, (2) strong evidence of MGPC bias toward the RGC fate and, (3) that knocking out 18 genes, of 100 tested, altered RGC regeneration kinetics. In a test of context specificity, 19 of 19 genes implicated in other regenerative paradigms were not required for RGC regeneration. Further, 13 of 14 genes shown to be required for regeneration following widespread retinal damage(8) were also dispensable for RGC regeneration. Intriguingly, knockout (KO) of *ascl1a* and *sox2* actually accelerated RGC regeneration kinetics.

We were particularly interested in *ascl1a* as overexpression of Ascl1 stimulates mammalian MG to produce new neurons(9,10). However, Ascl1-overexpressing mouse MG rarely divide, instead transdifferentiating directly into amacrine and bipolar retinal interneurons. New RGCs and photoreceptors, the cells most closely associated with retinal disease(11,12), are rarely observed. More recently, overexpression of Ascl1 and Atoh1 resulted in mammalian MG becoming “RGC-like” cells(13), showing cell fate can be modulated to promote the regeneration of disease-relevant cell types. This work highlights the need to better understand how proliferation and cell fate are controlled in reactive MG and MGPCs. In mechanistic analyses, we determined *ascl1a* KO in zebrafish had no effect on proliferation but enhanced MGPC bias toward the RGC fate. These findings demonstrate that the molecular regulation of retinal regeneration is context specific; i.e., divergent across paradigms due to the regenerative process being informed by, and actively adapting to, the nature of the retinal injury/cell loss. By extension, this suggests cell-specific regeneration paradigms will help to advance strategies for regenerating disease-relevant cell types selectively.

## Results

### A novel zebrafish RGC regeneration paradigm

To investigate factors regulating RGC regeneration, we established transgenic zebrafish enabling prodrug-inducible selective RGC ablation. In this “RGC:YFP-NTR2” line, a yellow fluorescent protein (YFP) reporter and improved bacterial nitroreductase (NTR 2.0)(14) enzyme are co-expressed exclusively in >95% of RGCs(15,16) (Fig. 1A; fig. S1A). This allows NTR-expressing RGCs to be ablated upon exposure to NTR prodrug substrates, such as metronidazole (Mtz)(14). To establish an RGC ablation paradigm amenable to large-scale screening, we exposed 5 days post-fertilization (dpf) RGC:YFP-NTR2 larvae to a range of Mtz concentrations and quantified YFP levels at 7 dpf using a fluorescence plate reader assay(17). Mtz treatments of ≥100 μM for 24 (Fig. 1B) and 48 hr (fig. S1B) were sufficient to reduce YFP expression to non-transgenic (non-Tg) control levels. Accordingly, all subsequent assays used a 24 hr 100 μM Mtz treatment to induce maximal RGC loss. To assess if reductions in YFP correlated to RGC loss, nuclei in the ganglion cell layer (GCL) were quantified in retinal sections of control and Mtz-treated larvae at 7 dpf. Mtz caused an ∼75% reduction in GCL nuclei (fig. S1C,D), with many remaining GCL nuclei likely being displaced amacrine cells(18).

**Fig. 1.**
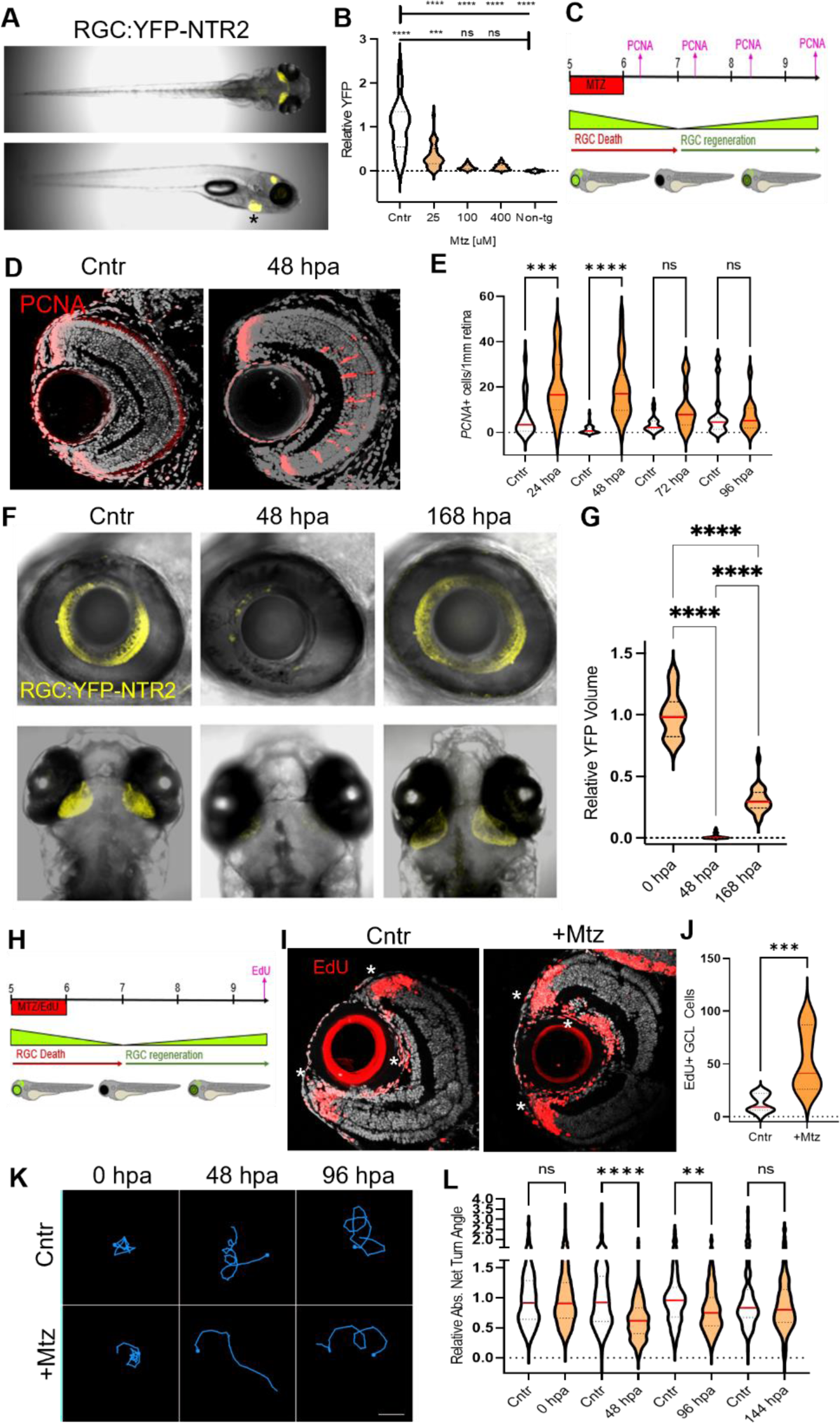
Characterization of RGC:YFP-NTR2 paradigm. (**A**) Whole fish view of RGC:YFP-NTR2 expression (*note, asterisk indicates reporter expression in the heart due to a “tracer” transgene, not YFP-NTR2 expression*). (**B**) Quantification of YFP following Mtz-induced RGC ablation at the indicated concentrations. (**C**) Experimental setup to analyze proliferative responses to RGC ablation. (**D**) Representative PCNA immunostaining (red cells) showing proliferation in INL cell clusters following RGC ablation (note, peripheral labeling is ongoing proliferation in CMZ). (**E**) Daily quantification of INL proliferative response (PCNA^+^ cells) following RGC ablation (24-96 hpa). (**F**) *In vivo* imaging of ablation and regeneration of RGC cell bodies in retina (top) and RGC axons in tectum (bottom). (**G**) Volumetric quantification of Mtz-induced RGC axonal loss and regeneration in the tectum observed by *in vivo* imaging (YFP signal). (**H**) Experimental setup to acutely lineage trace MG-derived progenitor cells with EdU upon RGC ablation (24 hr pulse, 96 hr chase). (**I**) Representative images of EdU immunostaining in control and Mtz-treated larvae (+Mtz; note, asterisks indicate EdU labeling of CMZ cells at periphery and endothelial cells near the optic nerve layer that were not included in quantification). (**J**) Quantification of EdU^+^ cells in the GCL. (**K**) Representative traces of visual response to lights off stimulus in control and Mtz-treated larvae at 0, 48, and 96 hpa.(**L**) Quantification of visual responses to lights off stimulus of control and Mtz-treated larvae (0-144 hpa). All white violin plots are unablated control larvae (Cntr), all orange violin plots are RGC ablated larvae (+Mtz). Asterisks in plots indicate p-value range: *<0.05, **<0.01, ***<0.001, <0.0001. Abbreviations not previously defined: not statistically significant (ns).

Retinal damage is known to induce Müller glia (MG) to dedifferentiate to a stem-like state and divide to produce a MG-derived progenitor cell (MGPC). MGPCs are transit amplifying neural progenitor cells which divide further and differentiate into new retinal neurons(19,20). Accordingly, we asked if RGC loss induced MG/MGPC proliferation. Following Mtz-induced ablation at 5 dpf, RGC:YFP-NTR2 larvae were collected daily until 9 dpf (24 - 96 hours post-ablation, hpa) and processed for proliferative cell nuclear antigen (PCNA) immunostaining (Fig. 1C). Mtz treatment led to PNCA staining in closely associated chains of cells spanning the inner nuclear layer (INL), consistent with proliferation of MG and MGPCs(20) (Fig. 1D). Co-immunostaining assays performed at 1 dpa demonstrated colocalization of PCNA and glutamine synthetase (GS), a marker of MG cells (fig. S1E). Quantification of PCNA-positive cells showed increased proliferation at 24 and 48 hpa following Mtz treatment (Fig. 1E). Tests in juvenile stage fish (∼6 week old) showed similarly robust PCNA staining in the INL following Mtz-induced RGC ablation (fig. S1F). Unfortunately, variable transgene expression downregulation shortly thereafter made adult stage studies impractical.

To test if RGCs regenerate following ablation, Mtz-treated RGC:YFP-NTR2 larvae were allowed to recover until 11 dpf (168 hpa, Fig. 1E). Intravital confocal time series images were collected pre-Mtz treatment at 5 dpf (Cntr, 0 hpa), 48 hpa (7 dpf), and 6 days post-fertilization (dpa; 11 dpf). Confocal z-stack projections show clear evidence of RGC loss and regeneration in the Mtz-treated group (Fig. 1F). Quantification of YFP volumes in the tectum, corresponding to RGC axons, showed a ∼99% reduction at 48 hpa and a return to ∼30% of pre-ablation control levels at 6 dpa (Fig. 1G).

Studies using the NTR/Mtz cell ablation system provided initial evidence of “fate biased” retinal regeneration, MGPCs preferentially gave rise to lost cell types following selective amacrine cell and cone photoreceptor ablation(5,6,21). Fate-biased regeneration has recently been observed in light damage (LD) and N-methyl-D-aspartate (NMDA) excitotoxicity models as well(4). To investigate MGPC fate choices following RGC ablation, ethynyl-2’-deoxyuridine (EdU) was pulsed with Mtz from 0-24 hpa (5-6 dpf) to label proliferating cells and then chased to 96 hpa (9 dpf) to assess MGPC differentiation. Immunostaining revealed unablated controls had EdU-positive cells in the ciliary margin zone (CMZ) and the endothelial layer surrounding the lens, but rarely any EdU-positive cells in the GCL, INL, or outer nuclear layer (ONL; Fig. 1I, Cntr). In contrast, Mtz-treated retinas showed prominent EdU staining in the GCL and rare EdU-positive cells in other retinal layers (Fig. 1I, +Mtz). Quantification showed a statistically significant increase of EdU staining in the GCL of Mtz-treated retinas (Fig. 1J). Together with the absence of EdU cells in other retinal layers at this time point in both Cntr and +Mtz images, this data suggests targeted RGC ablation induced a fate biased regenerative response, similar to selective amacrine and photoreceptor ablation paradigms(5–7).

We next assessed whether RGC loss and regeneration were correlated to changes in a visually-driven behavior. Upon loss of light, zebrafish larvae exhibit a phototaxis response characterized by an initial area-restricted search (helical turning) followed by a roaming search pattern(22–24). Only the initial area-restricted search requires vision(25). We therefore used this assay to test if RGC ablated larvae show altered initial responses to darkness onset. RGC ablation significantly impacted darkness-induced behaviors relative to controls at 48 hpa and 96 hpa, before returning to baseline levels at 6 dpa (Fig. 1K-L). The observed kinetics of RGC ablation and regeneration (e.g., YFP levels; Fig. 1F,G) therefore parallel the observed timing of lost and re-established visual performance in this assay.

### Transcriptomic analysis of RGC regeneration

To gain mechanistic insights, we profiled transcriptional changes associated with RGC loss and regeneration using time resolved single-cell RNA sequencing (scRNA-seq). Whole eye samples were collected from both RGC ablated and control larvae at 12, 24, 48, and 72 hpa (Fig. 2A). A total of 133,104 cells were profiled. UMAP clustering identified all major cell classes of the eye (Fig. 2B), as confirmed by known markers (fig. S2). A total of 4,949 differentially expressed genes (DEGs) were identified (minimum FC>1.18, adjusted p-value <0.05, Fig. 2B, table S1). DEGs were enriched in activated MG (1,386) and dying RGCs (174) at 12 hpa, activated MG (478) at 24 hpa, retinal progenitor cells (MGPCs and CMZ cells; 317) at 48 hpa, and MG (206) and RGCs (223) at 72 hpa (Fig. 2C).

**Fig. 2.**
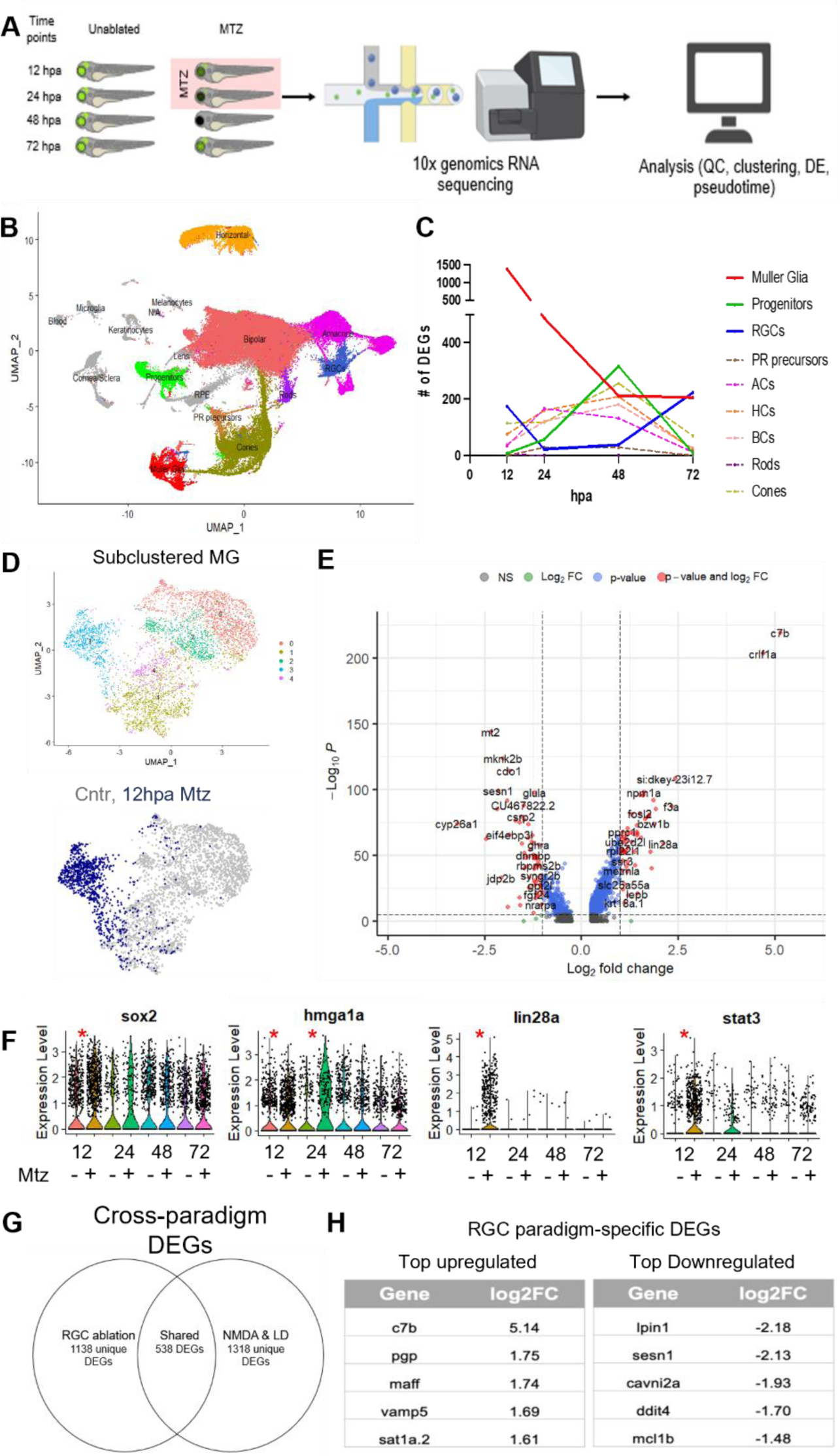
Transcriptomic profiling of biased RGC regeneration. (**A**) Experimental setup for scRNA-seq time series (12, 24, 48, and 72 hpa) of unablated control and RGC ablated larvae (+Mtz). (**B**) UMAP of annotated eye cells from all timepoints collected. (**C**) Total DEGs identified at each timepoint for MG, progenitor cells, RGCs and other major retinal neuron classes. (**D**) Subclustering revealed a reactive MG subcluster made up predominantly of 12 hpa cells. (**E**) Volcano plot of DEGs in 12 hpa MG subcluster. (**F**) Select genes among early timepoint DEGs in MG following RGC ablation (asterisks indicate statistical significance). (**G**) Cross-paradigm comparison of MG DEGs from RGC ablation data and a pooled NDMA/LD dataset. (**H**) Top RGC paradigm-specific upregulated and downregulated DEGs in the RGC ablation dataset and pooled NMDA/LD datasets. Abbreviations not previously defined: amacrine cells (ACs), bipolar cells (BCs), horizontal cells (HCs), photoreceptor (PR), quality control (QC). All gene abbreviations are per current Zebrafish Information Network (ZFIN) nomenclature.

To assess specific gene changes in MG following RGC ablation, the MG cluster (Fig. 2B, red) was isolated and subclustered. Five subclusters were identified, including a subcluster (#3) almost entirely composed of 12 hpa MG cells (Fig. 2D). A volcano plot of 12 hpa MG DEGs shows two highly induced genes observed in MG single cell data other models of retinal regeneration, *c7b*(21,26) and *crlf1a*(27) (Fig. 2E). Other MG activation markers enriched in 12 hpa MG included *sox2*, *hmga1a*, *lin28a*, and *stat3* (Fig. 2F). Interestingly, *ascl1a*, a gene previously shown to be upregulated and required for MG activation in retinal tissue damage paradigms(8,28,29), was absent from the list of DEGs enriched in activated MG following RGC ablation.

Next, we compared MG DEGs in a related dataset. Hoang et. al performed scRNA-seq of adult zebrafish retinas following widespread retinal cell loss upon light damage (LD) or injection of the NMDA(29). Pooling MG datasets from NMDA and LD paradigms identified 1,856 DEGs, versus a total of 1,678 MG DEGs identified in our data. Comparisons across datasets revealed 538 shared DEGs (32%), while 1,138 were unique to the RGC paradigm (Fig. 2G, table S2). This proportion of paradigm-specific and shared DEGs parallels a bulk RNA comparison of two selective cell ablation paradigms(21). Upregulated MG DEGs unique to our dataset included *c7b*, *pgp*, *maff*, *vamp5*, and *sat1a.2*, while unique downregulated genes included *lpin1*, *sesn1*, *cavin2a*, *ddit4* and *mcl1b* (Fig. 2H).

### Pseudotime trajectory identifies RGC ablation-induced genes across regeneration

We next employed pseudotime analyses to assess gene changes along a MG>Progenitor>RGC trajectory, akin to related studies(4,13,29,30). We isolated MG, progenitor cells (both MGPCs and CMZ cells) and RGCs from control and ablated samples and inferred a common pseudotime trajectory (Fig. 3A). Analysis of cell density along the trajectory showed enrichment of ablated sample cells early during MG activation, less differences in the middle where progenitor cells were enriched, and enrichment for control sample cells later (Fig. 3B). 1,829 DEGs were identified along pseudotime which segregated into 11 expression patterns (fig. S3A, table S3). Genes upregulated in Clusters 1 and 2 genes were associated with inflammation (*c7b*, *crlf1a*), MG activation (*gfap*, *lin28a*) and DNA damage (*gadd45ab*). Genes upregulated in Cluster 3 included markers of MGPCs (*pax6a*, *foxn4*, *sox2*) and RGC precursors (*atoh7*, *pou4f2*). Cluster 4-7 genes were upregulated progressively later in the trajectory, including neurogenic factors (*thrb*, *neurod1/4/6b*, *otx2b*) and genes associated with neuronal differentiation (*gap43, alcamb*). Cluster 8-11 genes were downregulated progressively later and included neurogenic factor (*zfhx3*) and mature neuronal markers (*elavl4)*. Gene ontology analysis on each cluster identified multiple developmentally important signaling pathways including MAP kinase, JAK/STAT, Wnt, BMP, and Hedgehog (fig. S3B).

**Fig. 3.**
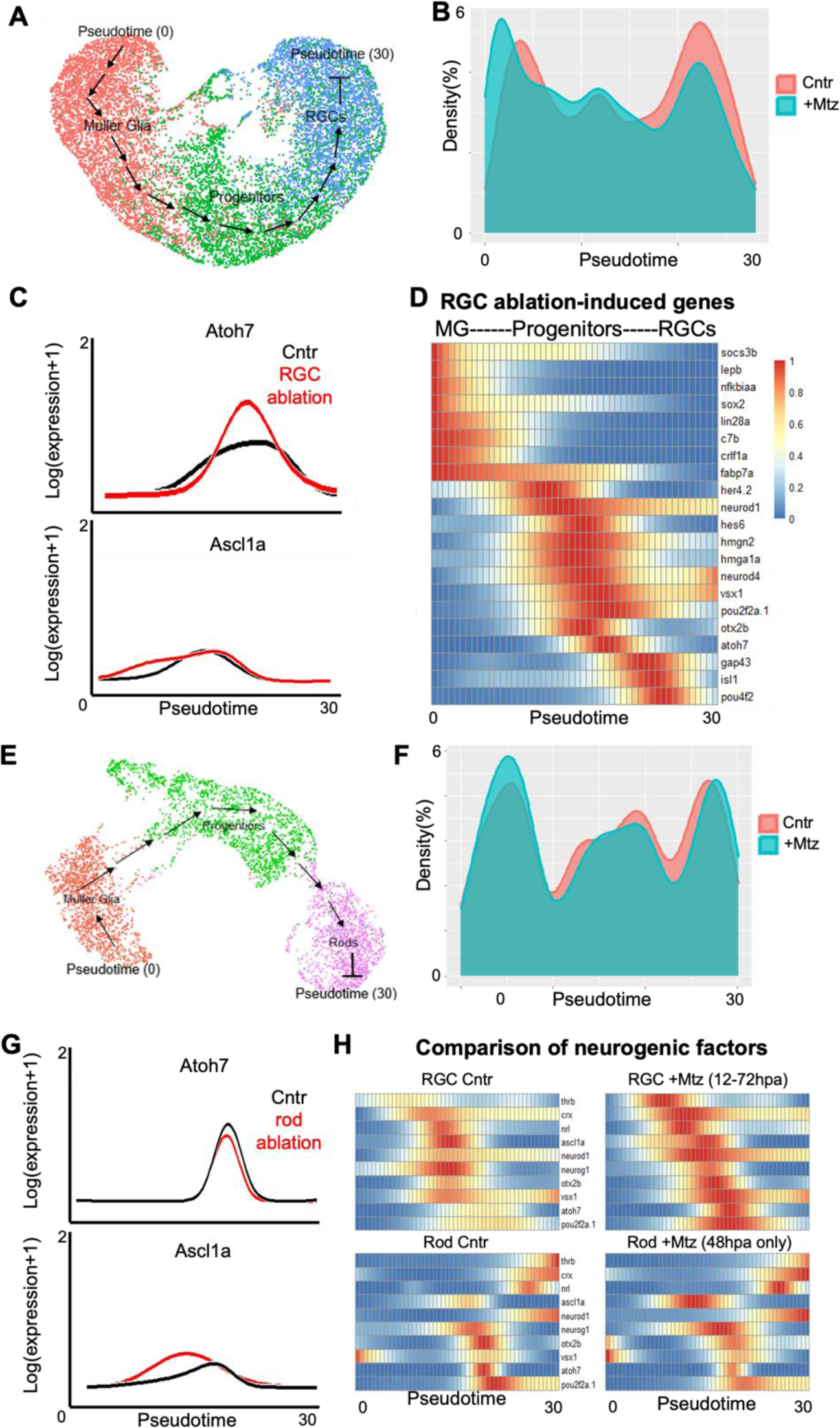
Pseudotime analysis of RGC and Rod regeneration. (**A**) UMAP used to create pseudotime MG>Progenitors>RGCs trajectory from 0-72 hpa data. (**B**) Density of unablated control (Cntr) and RGC ablated cells (+Mtz) along trajectory. (**C**) Expression of *atoh7* and *ascl1a* in Cntr and RGC ablated cells along trajectory. (**D**) Heatmap of select DEGs upregulated during RGC loss and regeneration. (**E**) UMAP used to create pseudotime MG>Progenitors>RGCs trajectory from 48 hpa data. (**F**) Density of unablated control (Cntr) and rod cell ablated (+Mtz) along trajectory. (**G**) Expression of *atoh7* and *ascl1a* in Cntr and rod ablated cells along trajectory. (**H**) Comparison of changes in expression of select neurogenic factors between RGC ablation (top) or rod cell ablation (bottom) paradigms along corresponding pseudotime trajectories. All gene abbreviations are per current Zebrafish Information Network (ZFIN) nomenclature.

We were particularly interested in expression patterns of transcription factors *atoh7* and *ascl1a*. During development, *atoh7* specifies the RGC lineage(31). As mentioned above, *ascl1a* has been shown to be required for retinal tissue regeneration(8). Following RGC ablation, *atoh7* was strongly induced in the progenitor phase while *ascl1a* exhibited a minor increase earlier during the MG activation phase (Fig. 3C). A subset of the most significantly induced genes along the trajectory (Fig. 3D) included factors associated with: (1) MG activation (*lepb*, *nfkbiaa*), (2) MGPCs (*sox2*, *fabp7a*, *her4.2*), (3) neurogenesis (*neurod1*, *neurod4*, *hmgn2*, *vsx1*),(4) specification of RGCs (*atoh7*) and other early developmental cell fates (*pou2f2a*), and lastly (5) RGC differentiation and maturation (*isl1*, *pou4f2,* and *gap43*)(32).

To assess context specificity of the RGC regeneration transcriptomic signature, we performed scRNA-seq using a published NTR/Mtz-based rod photoreceptor ablation model(17). 5 dpf larvae were treated ±Mtz for 24 hrs and whole eye samples collected at 48 hpa. Pseudotime analysis was used to construct a MG>Progenitor>Rod trajectory (Fig. 3E-F). In contrast to the RGC paradigm, rod ablation led to a decrease in *atoh7*, and a stronger increase in *ascl1a* expression (Fig. 3G). Comparing neurogenic gene profiles showed obvious differences related to neuronal cell type of each trajectory (Fig. 3H; RGC Cntr vs. Rod Cntr), and paradigm-specific changes following induction of cell loss. RGC ablation induced relatively higher levels of *atoh7*, *pou2f2a, thrb*, and *otx2b* while rod loss led to higher relative levels of *ascl1a* and *neurog1*,(Fig. 3H).

### Reverse genetics screen: knockout of *ascl1a* enhances RGC regeneration

To identify regulators of RGC regeneration, we performed a “crispant” genetic screen to evaluate the effects of knocking out 100 genes on RGC regeneration kinetics. This CRISPR/Cas9 multi-guide RNA approach creates biallelic mutations in targeted genes in F0 generation zebrafish embryos. Candidate genes were chosen from our scRNA-seq dataset and prior published reports(29). Plate reader-based quantification determined that ∼32% of RGCs had regenerated in non-mutated control larvae by 72 hpa (Fig. 4A,B). Compared to ablated controls, KO of eleven increased RGC replacement kinetics (*ascl1a*, *neurog1*, *max*, *sox2*, *olig2*, *lepb*, *tubb4b*, *nfasca*, *tnfrsf11b*, *fosab*, and *mink1*) while KO of seven inhibited RGC regeneration (*cry2b*, *slc1a2b*, *mlc1*, *arntl2b*, *hspd90bl*, *atf6* and *ctnnb1*; Fig. 4C, table S3).

**Fig. 4.**
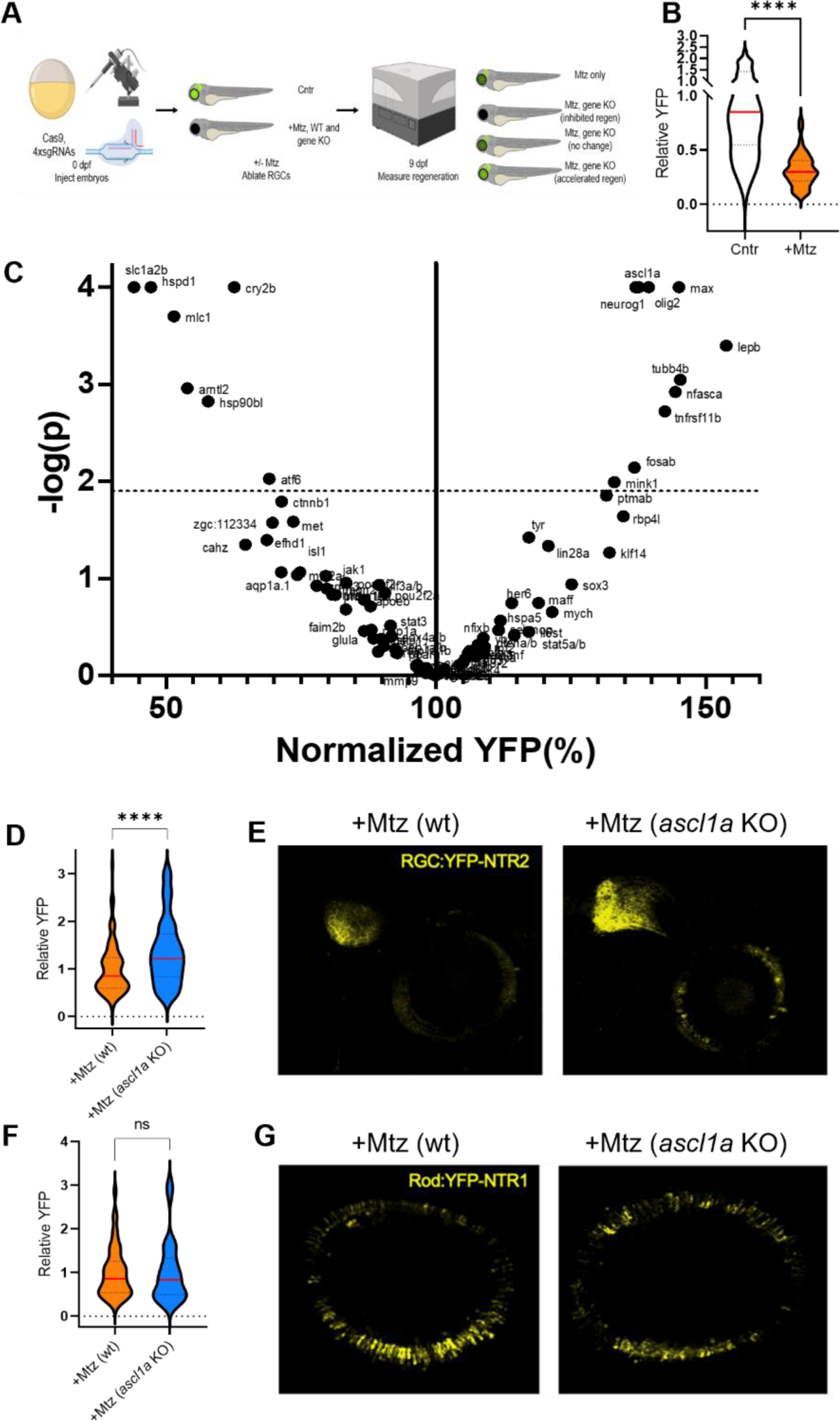
Reverse genetics screen identifies mediators of RGC regeneration. (**A**) Experimental setup for reverse genetics CRISPR/Cas9-enabled “crispant” knockout (KO) screen for genes that regulate RGC regeneration. (**B**) Quantification of YFP signal from RGC:YFP-NTR2 samples at 96 hpa for unablated control (Cntr) and RGC ablated control larvae, the timepoint chosen for assessing changes in RGC replacement kinetics. (**C**) Volcano plot of normalized YFP levels at 96 hpa for all Mtz-treated RGC:YFP-NTR2 crispant KO larvae (dashed line indicates adjusted p-value cutoff of 0.01, minimum sample size n>8, median sample size n=16). (**D**) Quantification of YFP signal in RGC ablated control (+Mtz) and RGC ablated *ascl1a* crispant larvae (+Mtz, *ascl1a* KO) at 96 hpa. (**E**) Representative images of RGC regeneration in +Mtz and +Mtz *ascl1a* KO larvae at 168 hpa. (**F**) Quantification of YFP signal from Rod:YFP-NTR1 samples at 96 hpa for Rod cell ablated control (+Mtz) and Rod cell ablated *ascl1a* crispant larvae (+Mtz, *ascl1a* KO). (**G**) Representative images of Rod cell regeneration in +Mtz control and +Mtz, *ascl1a* KO larvae at 168 hpa. All gene abbreviations are per current Zebrafish Information Network (ZFIN) nomenclature.

Intriguingly, KO of several genes known/implicated as being required for retinal regeneration following widespread tissue damage, e.g., *ascl1a*(8), *sox2*(33), *olig2*(34), and *lepb*(35), resulted in accelerated RGC regeneration kinetics. In contrast, KO of *mmp9* had no effect on RGC regeneration(4), despite roles in regulating proliferation following LD and promoting INL and GCL fates in both LD and NMDA paradigms(4) (Table S4). We were particularly intrigued by the *ascl1a* KO result as overexpression of Ascl1 in mouse MG has been used to stimulate a nascent regenerative response in the injured mouse retina(13,30). Follow up plate reader assays (Fig. 4D) as well as *in vivo* imaging (Fig. 4E) confirmed primary screen results, showing *ascl1a* KO enhanced RGC regeneration. Additional controls showed *ascl1a* KO of was highly efficient and had no effect on RGC development, or Mtz-induced RGC death (fig. S4). To assess specificity, we tested the effect of *ascl1a* KO on rod photoreceptor regeneration kinetics and saw no change by either plate reader assay (Fig. 4F) or intravital imaging (Fig. 4G).

Consistent with this, functional tests of the 33 genes known/implicated in regeneration in the context of widespread damage (LD, NMDA, puncture wounds) found that most genes affected only a single retinal regenerative paradigm or had opposing effects in different models, such as *ascl1a* and *sox2* (table S4).

### Ascl1a knockout biases toward early progenitor cell fates in MGPCs

To examine how *ascl1a* KO enhanced RGC regeneration, we performed a multiomic snRNA-seq/ATAC-seq analysis on wt and *ascl1a* KO retinas at 0 and 24 hpa (Fig. 5A), profiling a total of 53,889 cells. UMAP clustering of an integrated dataset identified all major cell classes (fig. S5). Analysis of differential chromatin accessibility in 24 hpa retinal progenitor cells identified significantly increased accessibility peaks for *atoh7* in *ascl1a* KO cells (Fig. 5B). We next analyzed scATAC-seq data in MG and progenitor clusters for differentially accessible transcription factor binding motifs. Motifs with increased accessibility in *ascl1a* KO MG included Atoh1, Olig1, Nr2f2, Neurod2, and Otx1, while decreased motifs included Nrf1 and Ascl1 (Fig. 5C). For *ascl1a* KO progenitor cells, motifs with increased accessibility included E2f2, Six3, Sox4/10 and Vsx1/2. Motifs with decreased accessibility included Pitx1, Otx2, Thrb, Stat1 and Ascl1 (Fig. 5D).

**Fig. 5.**
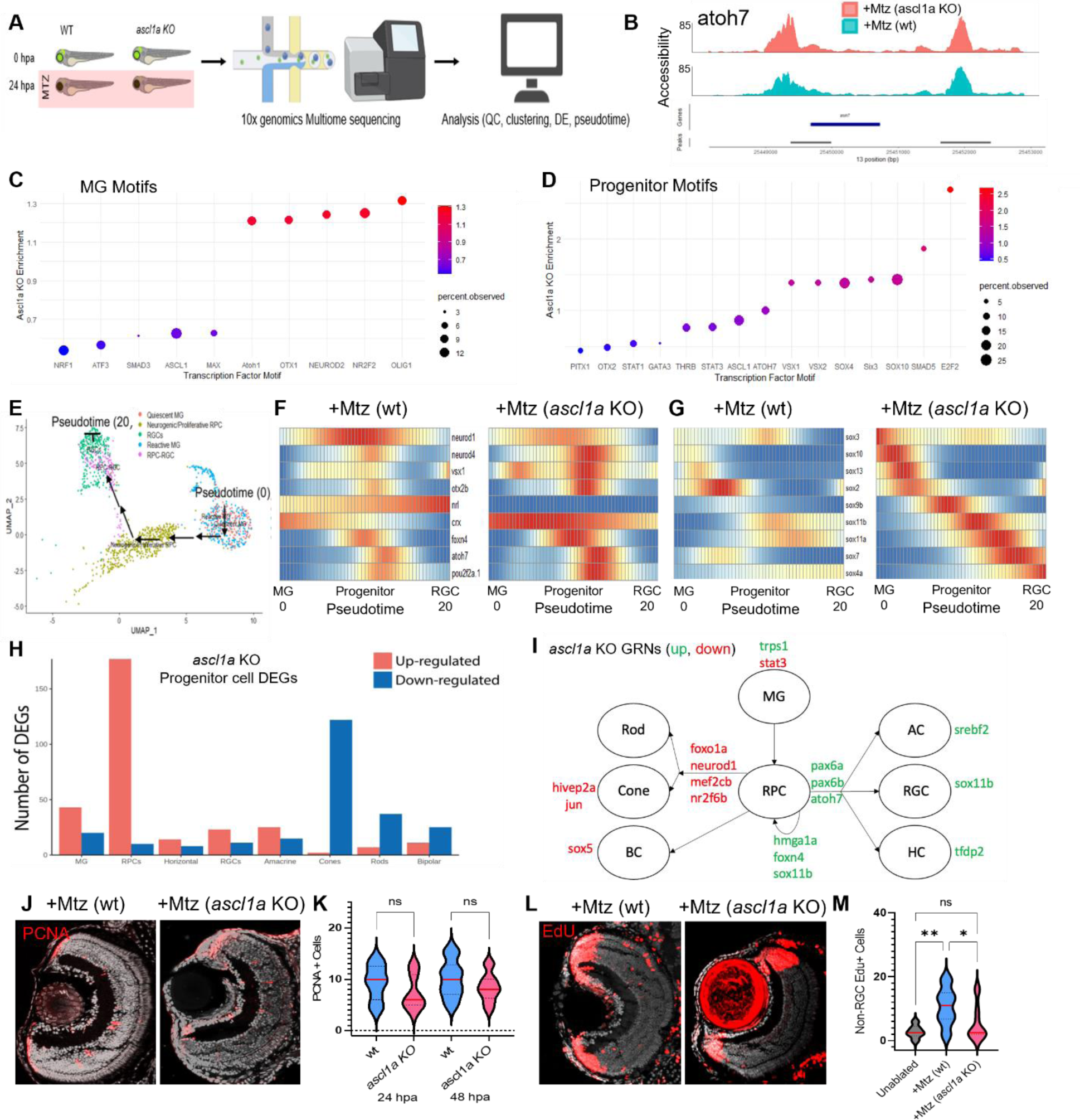
Ascl1a KO accelerates RGC regeneration kinetics by enhancing RGC fate bias. (**A**) Experimental setup for multiome sequencing (combined snRNA-seq and ATAC-seq) at 0 and 24 hpa of wildtype (wt) control and *ascl1a* KO crispant larvae. (**B**) DNA accessibility comparison between wt and ascl1a KO larvae at the *atoh7* locus following RGC ablation (+Mtz). (**C**) Transcription factor motif accessibility enrichment in MG cells of *ascl1a* KO crispants. (**D**) Transcription factor motif accessibility enrichment in progenitor cells of *ascl1a* KO crispants. (**E**) UMAP used to create pseudotime trajectory from quiescent MG to RGCs. (**F**,**G**) Heatmap of select pro-neural and sox TFs expression changes across pseudotime for wt and *ascl1a* KO larvae at 24 hpa following RGC ablation. (**H**) DEGs in progenitor cells of *ascl1a* KO larvae associated with known retinal cell lineages. (**I**) Derived gene regulatory networks (GRNs) altered in *ascl1a* KO progenitors based on changes in expression of transcription factors known to regulate DEGs in (H). (**J**) Representative PCNA immunostaining in RGC ablated control (+Mtz) and RGC ablated ascl1a crispant (+Mtz, *ascl1a* KO) retinas. (**K**) Quantification of PCNA^+^ cells following RGC ablation at 24 and 48 hpa in wt and *ascl1a* KO retinas. (**L**) Representative EdU immunostaining images of “secondary” EdU lineage traced cells (0-48 h EdU pulse, 6-day chase) in RGC ablated wt and *ascl1a* KO retinas. (**M**) Quantification of the EdU^+^ retinal cells in the ONL and INL (excluding CMZ and GCL) in unablated controls and RGC ablated wt and *ascl1a* KO retinas at 6 dpa. All gene abbreviations are per current Zebrafish Information Network (ZFIN) nomenclature.

Next, we produced a MG>Progenitor>RGC pseudotime trajectory with annotation of new subclusters made possible by chromatin data (Fig. 5E), identifying 269 significantly differentially regulated factors (table S5). As above, we compared relative levels of neurogenic (as well as Sox family) TFs between wt and *ascl1a* KO retinas. Most neurogenic genes were co-expressed in progenitors, suggesting induction of a “transitional” progenitor cell state(36,37). Neurogenic TFs upregulated in *ascl1a* KO progenitor cells included: *crx*, *foxn4*, *neurod4*, *vsx1*, *otx2b*, *pou2f2a*, and *atoh7* – and all Sox genes evaluated, except *sox2*. Strong induction of *sox11a/b* and *sox4a* – both implicated in RGC development in mice(38) – suggests enhanced RGC fate bias in ascl1a KOs. Conversely, *neurod1* and *nrl* showed reduced levels of expression in *ascl1* KOs (Fig. 5G), suggesting reduced production of photoreceptors.

To further explore potential changes in cell fate, we examined changes in established lineage-promoting genes between wt and *ascl1a* KO progenitors. The data showed *ascl1a* KO increased the relative numbers of genes associated with RGC, horizontal cell and amacrine cell lineages, and decreased the number of genes associated with rod, cone and bipolar cell fates (Fig. 5H). To construct GRNs, we identified TFs predicted to underlie changes in *ascl1a* KO progenitor cells. TFs associated with downregulation of photoreceptor lineage genes included *neurod1*, *nr2f6b*, *foxo1a*, and *mef2cb*. TFs associated with upregulation of RGC, horizontal cell and amacrine cell lineage genes included *pax6a/b* and *atoh7* (Fig. 5I, table S6).

Lastly, to investigate whether disrupting *ascl1a* altered proliferation and/or cell fate bias during RGC regeneration, we used PCNA immunolabeling and EdU lineage tracing. Interestingly, *ascl1a* KO retinas exhibited no change in PCNA cell numbers relative to wt controls (Fig. 5J,K). We next assessed MGPC production of cells other than RGCs as a measure of relative fate bias. Similar to a NTR-based amacrine cell regeneration paradigm, acute ablation of nearly all RGCs is predicted to subsequently cause low levels of cell loss in the INL and ONL(6). To track MGPCs responding to secondary cell loss, the EdU pulse was extended by 24 hrs (0-48 hpa) and chased until 7 dpa (12 dpf). The data showed wt MGPCs gave rise to a small but statistically significant number of cells in INL and ONL following RGC ablation (Fig. 5L,M). In contrast, no significant differences were observed between unablated controls and *ascl1a* KO retinas, consistent with reduced levels of non-RGC production (Fig. 5L,M). Combined with our GRN analysis, these results suggest the mechanism by which *ascl1a* KO accelerated RGC replacement kinetics was by enhancing the level of RGC fate bias exhibited by MGPCs.

## Discussion

Our results demonstrate plasticity and context specificity in how retinal regeneration is regulated; *ascl1a*-independent mechanisms of MG activation and a preponderance of paradigm-specific gene effects, respectively. These findings echo emerging evidence of divergence in how retinal regeneration is regulated across paradigms, suggesting regenerative processes are informed by and adapt to the nature of the injury incurred(4,21). Where this information originates (dying and/or surviving cells), how it is transmitted, and how it controls proliferative and cell fate decisions are areas of future research. In support of this concept, divergent gene expression was evident in cross-paradigm single cell omics analyses; ∼70% of MG DEGs were paradigm-specific rather than shared across RGC ablation and LD/NMDA models. Similarly, a comparison of two selective retinal cell ablation models showed paradigm-specific gene changes also predominated(21).

To test for context-specific gene function, 33 previously identified known/implicated regulators of retinal regeneration were screened for effects on RGC regeneration. The majority of genes required for (13 of 14) or implicated in (19 of 19) regeneration in the context of widespread retinal damage had either no statistically significant effect (26 genes) or accelerated (7 genes) RGC regeneration kinetics. Only 2 crispants showed concordant effects across paradigms: *hspd1* KO inhibiting(39), and *pparg* having no effect(21) on retinal regeneration. We also saw no effects on RGC regeneration upon targeting JAK/STAT signaling (*stat3*, *stat5a/b*, *jak1*), which differs from results in retinal damage(40–43) and selective rod cell ablation models(21). Pseudotime analysis implicated *stat1b* and *stat2* in RGC regeneration, similar to bulk RNA analysis of a bipolar cell ablation model(21). These results highlight the need for functional gene testing across different regenerative models to account for paradigm-specific effects.

We identified 18 genes whose disruption altered RGC regeneration kinetics. Among the 7 crispants that inhibited RGC replacement, i.e., RGC regeneration-promoting genes, most have no known role in regeneration, except *hspd1*(39) (above) and *hsp90bl* (i.e., effects on senescence during muscle regeneration(44)). Among 11 crispants that accelerated RGC regeneration kinetics, *ascl1a*, *sox2*, *olig2*, and *lepb*, stood out as genes previously shown/implicated as being required for retinal regeneration in the context of widespread damage(8,33–35). The effect of *sox2* disruption may be dosage dependent, with low expression levels allowing proper eye morphogenesis and promoting RGC differentiation(45). How disrupting *lepb* effects RGC regeneration is less clear as it is highly upregulated across tissue- and cellular-level regenerative paradigms(35,46), including the RGC model. In the context of a retinal puncture wound, morpholino knockdown of the *lepb* (leptin) receptor (*lepr*) decreased proliferation of cells in the INL, but also reduced expression of *ascl1a* – the latter providing a potential connection between *lepb* and *ascl1a* KO results here. Our most intriguing result, was that disruption of *ascl1a* increased RGC regeneration kinetics.

We expected *ascl1a* KO to inhibit RGC regeneration kinetics due to *ascl1a* being required for MG proliferation following puncture wounding, ouabain toxicity, or light damage in zebrafish(8,28) (Table S4). In addition, forced expression of Ascl1 in mouse MG cells awakens latent regenerative potential in NMDA damaged retinas(13,30). However, new cells are generated largely by transdifferentiation of Ascl1-overexpressing MG rather than proliferation and the resulting cell types are limited largely to amacrine and bipolar cells. Recent studies have shown that co-expressing Ascl1 with other neurogenic factors – Atoh1(13) or Pou4f2, and Islet1(30)– in mouse MG leads to the production of RGC-like cells that fail to mature fully. Here, loss of *ascl1a* enhanced RGC fate bias during regeneration, which is in keeping with Ascl1-expressing RPCs rarely producing RGCs during development(47,48). Removing *ascl1a* from the pool of neurogenic factors co-expressed in “transitional” progenitor cells(36,37) may therefore increase RGC differentiation probability by decreasing non-RGC options. This could also account for the pro-RGC regenerative effects of knocking out *olig2*(49) and possibly *neurog1*. Constitutive expression Ascl1 in mouse MG cells may actively repress RGC differentiation, thus Ascl1-independent means of dedifferentiating MG and/or methods for downregulating Ascl1 after MG activation, may promote RGC maturation. Regarding Ascl1-independent dedifferentiation, several other neurogenic factors, including *sox2, sox10*, *neurod1*, and *neurog2*, are able to reprogram astroglia cells into neurons(50), as per overexpression of Ascl1 in mouse MG cells. Of these, *sox10* is upregulated early in *ascl1a* KOs (Fig. 5G).

Possible explanations for the prevalence of paradigm-specific gene effects we observed are differences in age (larvae vs. adults) and/or methods: gene disruption (crispants vs. morphants), retinal injury (cell ablation vs. LD/NMDA/puncture), and assay measure (cell replacement kinetics vs. proliferation). Future experimentation will be required to investigate the impact of these differences. Crispants robustly recapitulate stable mutant phenotypes(51), enabling large-scale screens of regeneration-associated genes in larvae(52) or adults(53). Most prior tests, however, involved injecting morpholinos into the eye and electroporation(54,55). Phenotypic discordance between morphant and crispant tests may therefore arise due to differential immune responses, off-target effects, and/or crispant mutant mRNA triggered genetic compensation(56). Genetic compensation could explain the degree of plasticity we observe, and should be investigated from both an experimental and therapeutic perspective, i.e., leveraged to promote regeneration. Finally, by assessing regeneration kinetics we were able to show retinal cell regeneration can be promoted independent of increases in MG/MGPC proliferation, likely explaining at least some of the discordance between our findings and prior reports focused largely on effects on MG/MGPC proliferation.

In summary, our findings suggest context-specific regulatory mechanisms govern each phase of the retinal regenerative process. However, mechanisms for reprogramming MG to a stem-like state and regulating MGPC proliferation and differentiation also appear to be inherently plastic, which has profound implications for regenerative therapeutics. Recent observations that MGPC proliferation rates are matched to the level of cell loss(4,57), support the use of regenerative paradigms for defining feedback mechanisms of proliferative control(58). Similarly, that fate bias may be a generalizable feature of regenerative processes(4) – versus a feature unique to selective cell ablation paradigms – opens new opportunities to enhance understanding of how retinal cell differentiation is controlled. Insights on these fronts could support strategies for selectively stimulating the regeneration of disease-relevant retinal neuron types.

## Acknowledgements

**Funding**

National Institutes of Health Grant NRSA Individual Predoctoral Fellowship, F31 EY032790-01 (KE)

National Institutes of Health grant T32EY007143-24 (KE)

Brightfocus Foundation National Glaucoma Research grant G2020315 (JSM)

National Institutes of Health grant R01EY026580 (JSM)

National Institutes of Health grant P30EY001765-45 (Wilmer Eye Institute)

National Institutes of Health grant P20GM144230 (EH)

National Science Foundation grant OIA-2242771 (EH)

National Institutes of Health grant T32-GM133369 (JH)

## Authors Contributions

K.E and J.S.M contributed to experimental design in all aspects of the study and prepared the manuscript. K.E. carried out *in vivo*, histological and sequencing experiments as well as the majority of computational analyses. P.L and J.Q contributed to computational analysis design and execution for gene regulatory networks. T.H, I.P and S.B aided in design and execution of sequencing experiments. J.H and E.H designed and carried out behavioral assays. A.V.S and D.F.A aided in design of transgenic constructs for NTR2 fish. A.C, Z.C, S.N, F.M, L.Z, D.W, C.R, G.G, P.M, A.M, T.M and B.R aided in *in vivo* and histological experiments. J.T aided in pseudotime analyses and pathway analyses. M.T.S aided in manuscript writing and editing.

## List of Supplementary Materials

Materials and Methods

Figs. S1 to S5

Tables S1 to S6

## Materials and Methods

### Zebrafish husbandry and transgenic lines

All studies were carried out in accordance with recommendations by the Office of Laboratory Animal Welfare (OLAW) for zebrafish studies and an approved Johns Hopkins University Animal Care and Use Committee animal protocol. All fish were maintained using established conditions at 28.5◦C with a 14:10 h light:dark cycle. All larvae used were given 1-phenyl 2-thiourea (PTU) beginning at 1 dpf to facilitate screening for transgenic expression prior to experiments.

### Transgenic Lines

The Rod:YFP-NTR1 ablation line (*Tg(rho:YFP Eco. NfsB)gmc500)* has been previously published(1–3). To enable bipartite transgenic targeting of RGCs (RGC:YFP-NTR2), we combined the previously published Tg(isl2b.3:Gal4)zc65 known to label ∼95% of RGCs when combined with UAS driver elements(4), with the NTR2.0 variant recently published-Tg(5xUAS:GAP-tagYFP-P2A-nfsB_Vv F70A/F108Y)jh513(5) along with a neuronally-restricted Gal4 driver line, Et(2xNRSE-fos:KalTA4)gmc617(6). For all experiments, larvae were screened at 5 dpf for strong YFP expression in expected regions and reduced off target expression. In the RGC:YFP-NTR2 fish, we occasionally observed off-target expression in the trigeminal neuroglia, fish with this expression pattern were not used in any quantitative assays.

### Metronidazole mediated cell ablation

Cell ablation mediated by metronidazole (Mtz) was performed by first anesthetizing 5 dpf RGC:YFP-NTR2 or Rod:YFP-NTR1 larvae with 1x MS-222 (tricaine) and screening for strong YFP expression in expected regions. Fish were then split into appropriate +/− Mtz groups and placed into 6-well plates at the proper Mtz concentration diluted in 5mL of E3/PTU media. Fresh 10 mM Mtz stocks were made for each experiment.

### ARQiv scans to measure cell development, loss and regeneration

To assess induced RGC development (at 5 dpf), RGC loss (at 7 dpf) and RGC or rod cell regeneration (at 9 dpf), fluorescence quantification assays were performed using the ARQiv system, as previously described (7).

To determine the minimum sample size required to detect hit genes in the reverse genetics screen, standard deviation was calculated from ARQiv readings of 96 ablated fish at 96 hpa (the endpoint for the screen). We then used a power calculator (ClinCalc available online based on a published technique(8)) to determine a minimum sample size of 8 would enable identification of a 30% effect size on regeneration in mutant fish.

### *In vivo* Confocal Microscopy to assess cell loss and regeneration

All intravital imaging applied previously detailed protocols(9). Confocal z-stacks encompassing all retinal/brain fluorescence in the sample (step size, 5 microns) were collected. Image analysis was performed using FiJi for basic image processing (i.e., ImageJ v1.49b; NIH) or Imaris (v7.6.5; Bitplane) for nuclei counting following creation of a surface for DAPI or volumetric quantification using local background-based volumetric rendering of YFP signals.

### Statistical analyses

Statistical tests performed for all quantification data were performed in GraphPad Prism 9. When more than two groups were compared, we performed one-way anova tests with Dunnett’s multiple comparisons correction and significance was considered an adjusted p-value of <0.05. For the CRISPR/Cas9 mutagenesis screen, we employed a one-way anova followed by the recommended two-stage false discovery rate correction of Benjamini, Krieger, and Yekutieli, that cutoff significance at an adjusted p-value of 0.01.

### Tissue preparation, immunohistochemistry and confocal microscopy

For immunohistochemistry, larval zebrafish were euthanized in 20x MS-222, fixed in 4% paraformaldehyde (PFA) for at least 4 hr, washed three times in 1x PBS (phosphate buffered saline; EMD Millipore) and placed in 30% sucrose for ∼1h. Samples were then mounted in cryogel embedding medium, frozen in liquid nitrogen then stored at −80⁰C until sectioned in the lateral plane at 10-14 μm thickness with a cryostat. Sliced sections were collected on standard microscope slides. For immunolabeling, slides were air dried at room temperature for ∼1 hr, rinsed in 1xPBS and then re-fixed with 4% PFA for 15 min. PBST rinses (1xPBS +0.1% Tween20, Fisher Scientific) were conducted to remove trace PFA followed by 5 min antigen retrieval with SDS (1% Sodium Dodecyl Sulfate; Fisher Scientific) in PBS. The blocking phase was performed with 3% goat serum in PBDT (1xPBS, 1% BSA, 1% DMSO, 0.1% TritonX-100) for 30 min and incubated with primary antibody/1% goat serum/PBDT overnight at 4°C. The next morning, slides were rinsed in PBST, stained with secondary antibody/PBDT ∼2 hours in a light protected humidity chamber and cover-slipped (22×50 mm, Fisher Scientific). PBST rinses removed unbound secondary antibody. Samples were protected with Vectashield + DAPI (Vector Laboratories) and cover-slipped (24×50 mm, Fisher Scientific).

For EdU lineage tracing experiments, screened larvae (+/− Mtz) were given 40 uL of 10 mM EdU (Thermo Fisher) for either 24h or 48h and collected at multiple timepoints following. Following standard sectioning and PBS washing following sectioning as above, and EdU detecting reaction was performing with manufacturers instruction (Click-iT™ EdU Cell Proliferation Kit for Imaging, Alexa Fluor™ 647 dye, Thermo Fisher). Following EdU reaction steps, slides that received additional antibodies received primary antibody staining as above.

Primary antibodies included: mouse anti-PCNA monoclonal antibody (1:500; Sigma Aldrich), mouse anti-glutamine synthetase (1:200; Millipore), and mouse anti-HuC/D (1:500; Invitrogen). Secondary antibodies included: anti-mouse Alexa Fluor 488 (1:500; Life Technologies), anti-mouse Alexa Fluor 647 (1:500; Life Technologies), anti-rabbit Alexa Fluor 594 (1:500; Life Technologies).

Images were collected with an Olympus FV1000 Confocal Microscope (405, 440, 488, 515, 559, and 635nm laser lines). Stacked confocal images were obtained using a 40x oil immersion objective with a 5 µm step size, 130 µm aperture, and 10 µm total depth. ∼5-6 sections were collected per retina centered around the optic nerve. Images (Olympus .OIB format) were analyzed using ImageJ. Manual cell counts were for PCNA and EdU+ cells were averaged for each group.

### CRISPR/Cas9 mediated targeting

CRISPR/Cas9 mediated redundant targeting injections were performed utilizing the published strategy and gRNA table by Wu et al(10). 4 oligos per target were ordered as DNA oligos, assembled with the general CRISPR tracr oligo, and then transcribed using pooled in vitro transcription (HiScribe T7 High Yield RNA Synthesis kit, New England BioLabs) and cleaned up with the NEB Monarch RNA Cleanup kit. A mixture of all four sgRNAs (1 ng in total) and Cas9 protein (2.5 µM, IDT) were injected into one-cell stage embryos. Each injection cycle, gRNAs for the tyrosinase gene were injected that allow for confirmation of injection efficacy by looking for reduced pigmentation in injected embryos at 2 dpf.

### PCR and qPCR to verify *ascl1a* knockout

Following ascl1a gRNA injections, validation of DNA cutting was performed by DNA extraction, PCR and 1% gel electrophoresis with the following primers (Fwd: CCGCGAACACGTTCCCAATGGA, Rev: TGACACTCGGGACCCGTGGTTT) using the genotyping method in the GeneWeld paper(11) To validate mRNA expression was reduced following knockout, 2 dpf embryos were processed for quantitative PCR using a published protocol(12). Briefly, extracted mRNA samples were reverse transcribed (Qiagen Omniscript RT kit, Qiagen) and stored at −20°C. Samples were run in triplicate using the BioRad iQ SYBR Green Supermix (BioRad) in iCycler IQ 96 well PCR plates (Bio-Rad) on a BioRad iCycler equipped with an iCycler iQ Detection System. The protocol consisted of three phases: 1) 92°C for 10:00 min, 2) 50x 92°C for 00:15 min, 60°C for 01:00 min, 3) 81x 55°c 95°c for 00:10 min. β-actin served as the house keeping gene and 2^−ΔΔCT^ method was used for normalization to ensure equal amounts of cDNA for comparisons. qPCR primers were designed using the online tool QuantPrime.

### Single cell sample preparation and RNA sequencing

For each sample eyes were collected from 20-30 larval fish (40-60 retinas) in sibling unablated or post injury transgenic fish (12/24/48/72 hpa for RGC: YFP-NTR2 and 48 hpa in Rod:YFP-NTR1). Retinal cells were processed through published protocols for 10x genomics as previously described(13). Library preparation was then performed according to 10x genomics protocols for the version 3.1 kit and sequencing was conducted through the Johns Hopkins Single & Transcriptomics Core on a NovaSeq at ∼500 million reads per library.

### Single-cell RNA sequencing analysis

Raw reads were mapped to the Danio rerio GRCz10 using Cell Ranger vX.X from 10x genomics. Aligned genomic reads were then read into the published Seurat pipeline (v4.3.0.1) and quality control was performed by removing any cells with <200 detected genes or 1000 UMIs, and genes detected in fewer than 3 cells per experiment. Clustering steps were performed using steps from the pbmc Seurat tutorial available online. Briefly, the top 2,000 variable genes were identified and used to identify principal components (PCs) of the data. The top 30 PCs were used to produce a UMAP and clusters were annotated with known zebrafish marker genes(13). Differentially expressed genes (DEGs) were identified using the FindAllMarkers function between each control and ablation timepoint in each retinal cell cluster (minimum log2 foldchange cutoff of 0.25).

### Pseudotime trajectory analyses

Slingshot (v2.4.0) was used to infer pseudotime trajectories following subsetting of datasets into clusters of interest (i.e. MG, RPCs and appropriate neurons depending on the dataset). MG were treated as the root cluster and then the getLineages and getCurves functions were used to produce trajectories. In each case, common trajectories were learned using two conditions – either ablated and control cells, or wildtype and mutant cells from each appropriate experiment. The slingPseudotime function was used to calculate pseudotime for each cell and measure expression of each gene in pseudotime bins. Last, tradeSeq (v1.10.0) was used to compare gene expression between conditions (i.e. at pseudotime bin 1-calculate expression in control and ablated cells, for example), yielding lists of differentially expressed factors.

### Single cell multiome sample preparation and sequencing

For each sample eyes were collected from 20-30 larval fish (40-60 retinas) and flash-frozen in dry ice for ∼15min before being transferred to a −800 C freezer for storage. Nuclei were extracted from frozen retinal tissues according to 10xMultiome ATAC + Gene Expression (GEX) protocol (CGOOO338). Briefly, frozen retinal tissues were lysed in ice-cold 500ml of 0.1X Lysis buffer using a pestle and incubated on ice for 6 min totally. Nuclei were centrifuged, washed 3 times and resuspended in 10xMultiome nuclei buffer at a concentration of ∼3000-5000 nuclei/ml. Nuclei (∼15k) then were loaded onto 10x Genomic Chromium Controller, with a target number of ∼10K nuclei per sample. RNA and ATAC libraries were prepared according to the 10xMultiome ATAC + Gene Expression protocol and subjected for Illumina NovaSeq sequencing at ∼500 million reads per library

### Single-cell multiome sequencing analysis

RNA expression data was processed as above. Peak calling from single nuclei ATAC-seq reads was performed using MACS2 in the ArchR package (v1.0.2). ATAC seq data was then processed using the pbmc scATAC-seq workflow with the Signac (v1.10.0) and Seurat (v4.3.0.1) packages for quality control, normalization and producing an integrated UMAP. Differential expression and accessibility was then calculated for both gene RNA expression and chromatin peak accessibility. Next, the ChromVar package (v1.18.0) was used to identify differentially accessible transcription factor motifs between wildtype and *ascl1a* mutant cells.

### Identification of marker genes and differentially expressed genes for gene regulatory networks

To identify the differentially expressed genes (DEGs) in the RPC cell group between the control and ascl1a KO 24pha injury samples, we employed the “findMarkers” function from the Seurat (v4.3.0.1) package. DEGs were defined using the criteria: logfc.threshold > 0.2, min.pct = 0.05, and p-value < 0.05. Marker genes for each neuronal cell type were identified using the “findAllMarkers” function in Seurat with the following parameters: adjusted p-value < 0.05, min.pct = 0.05, and logfc > 0.5.

### Gene regulatory network construction

Using the snRNA-seq data from control and injury samples, we inferred TF-target relationships using the Arboreto package(v0.1.6)(14) in Python. We obtained importance scores for each TF-target pair using the “grnboost2” function. We then filtered the TF-target pairs based on these importance scores, removing pairs with scores lower than the 95th quantile. Additionally, we calculated the Pearson correlation for each TF-target pair based on the cell-by-gene expression matrix. A TF-target pair was annotated as “positive” regulation if its correlation (cor) was > 0.03, and as “negative” regulation if its correlation (cor) was < −0.03. Any other TF-target relationships were discarded. Finally, we filtered the GRNs based on TF expression in RPCs. Any TF not expressed in RPCs was filtered out.

### Identifying neuron-biased TFs upon *ascl1a* knockout

From the DEGs analysis, we noted that gene alterations in ascl1a knockout RPCs favor MG, RPCs, HC, RGC, and AC, while opposing Cone, Rod, and BC. To delve deeper into which TFs correlate with these gene expression biases, we identify RPC-expressed, neuron-favored TF activators that target specific neuron genes (similar to a recent study(15)). With the GRNs and neuron markers, we determined a specificity score (p-value) for each TF using a hypergeometric test using the “phyper” function in R. In this analysis, the “population” is the overall target genes from the GRNs, the “sample” comprises the TF target genes, and the “successes” are the neuron marker genes listed as target genes in the GRNs.

**Fig. S1.**
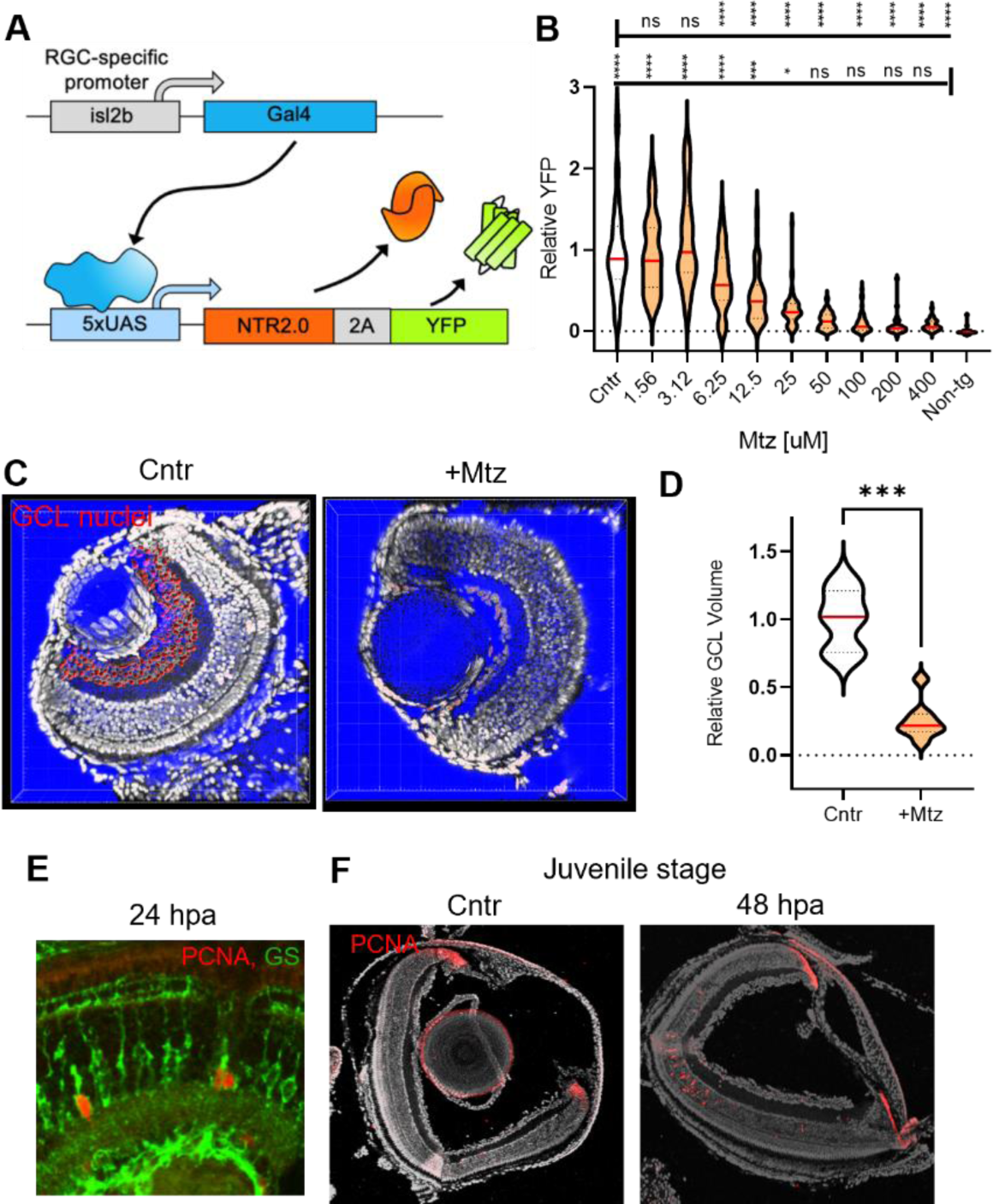
Additional characterization of the RGC:YFP-NTR2 model. (**A**) Schematic of transgenic strategy for RGC:YFP-NTR2 fish. (**B**) Plate-reader quantification of YFP-NTR2 signal following 48h Mtz treatment. (**C**) Imaris-processed representative images of sectioned retinas following Mtz ablation with GCL nuclei highlighted in red. (**D**) Quantification of GCL nuclei volume following ablation. (**E**) PCNA co-staining with *glutamine synthetase* (*gs*) following ablation. (**F**) PCNA staining in juvenile stage (6 week) RGC:YFP-NTR2 fish following ablation.

**Fig. S2.**
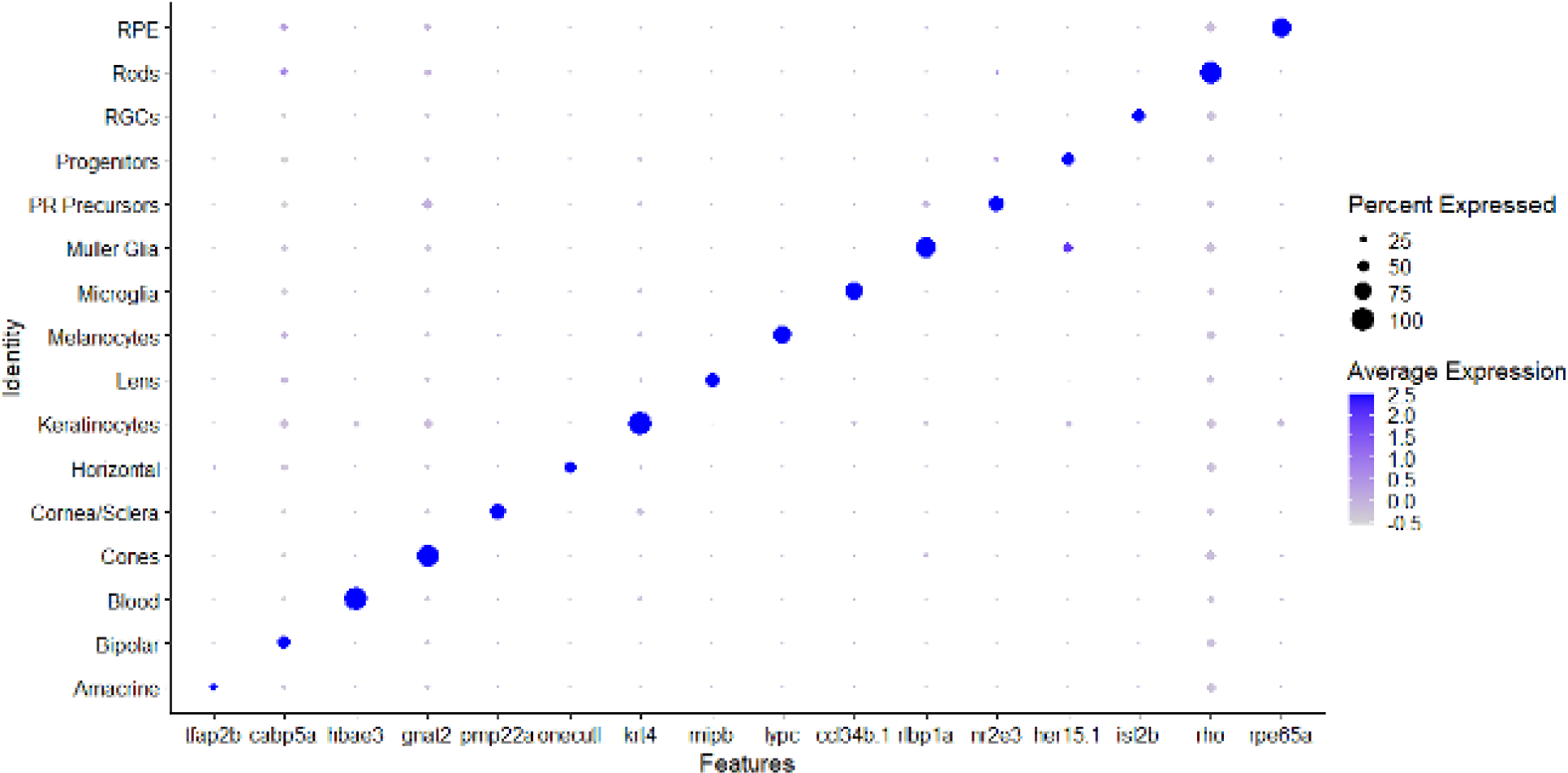
Identification of cell types using established marker genes. Marker genes used to establish cell types in scRNA data.

**Fig. S3.**
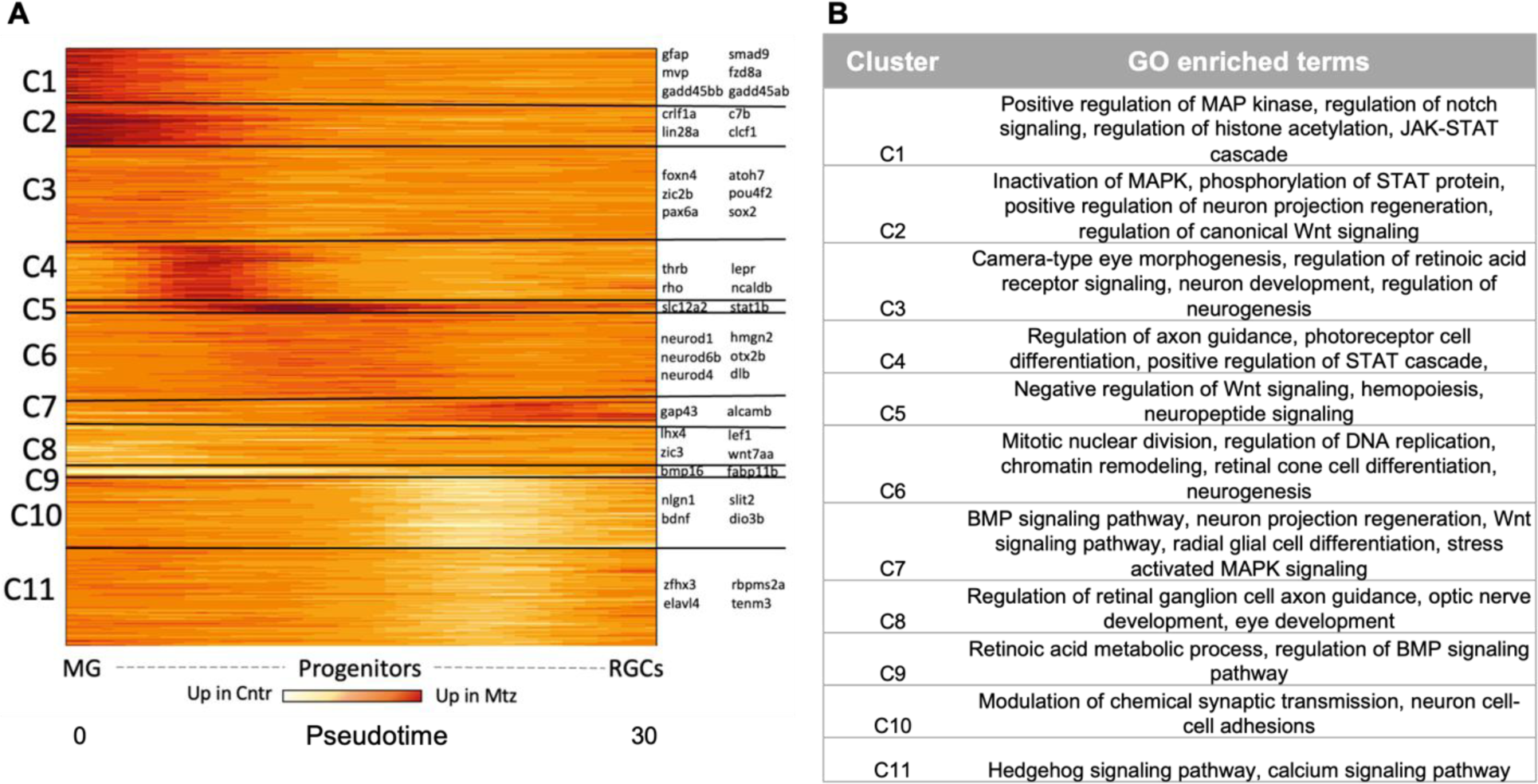
DEGs and associated GO cluster terms identified in MG>Progenitor>RGC pseudotime trajectory. (**A**) Subtractive heatmap (Mtz expr - Cntr expr) for 1,829 hit pseudotime DEGs clustered into 11 clusters (C1-C11), with example genes for each cluster to the right. (**B**) Gene ontology terms for significantly enriched pathways associated with each cluster.

**Fig. S4.**
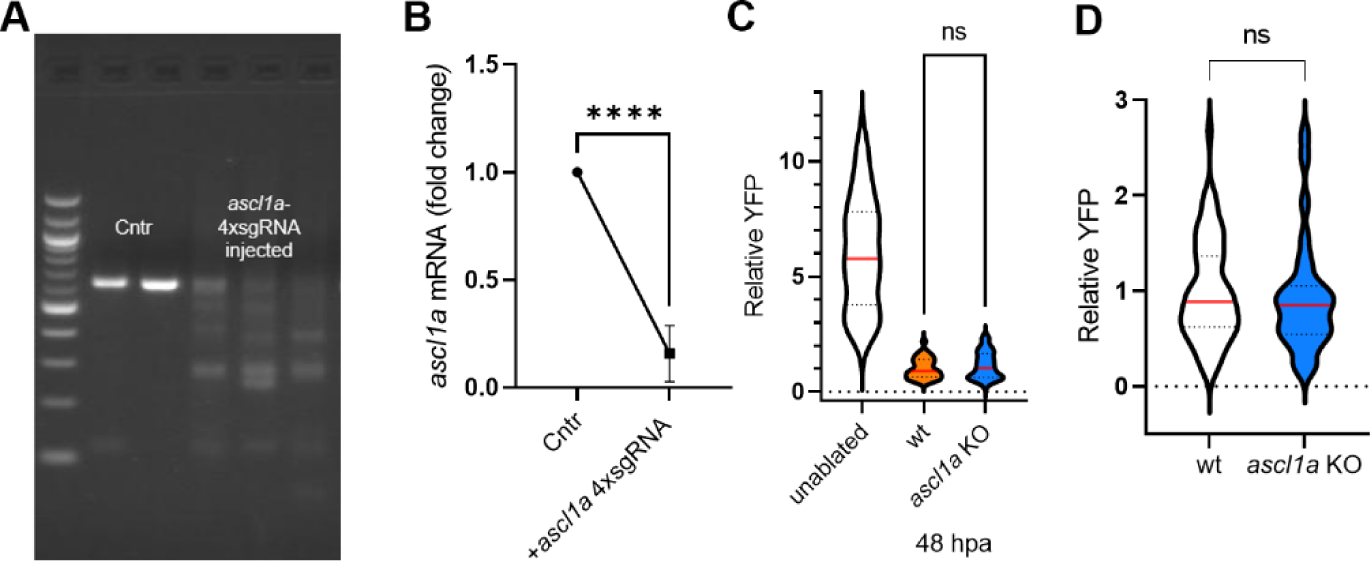
*Ascl1a* KO validation and effect on RGC development or Mtz-induced cell death. (**A**) PCR amplification of *ascl1a* coding sequencing following CRISPR/Cas9 knockout at 2 dpf. (**B**) mRNA expression of *ascl1a* following knockout at 2 dpf. (**C**) Comparison of Mtz-induced RGC ablation in ablated wt and ascl1a KO larvae at 48 hpa. (**D**) Comparison of RGC development at 5 dpf in wt and *ascl1a* KO RGC:YFP-NTR2.

**Fig. S5.**
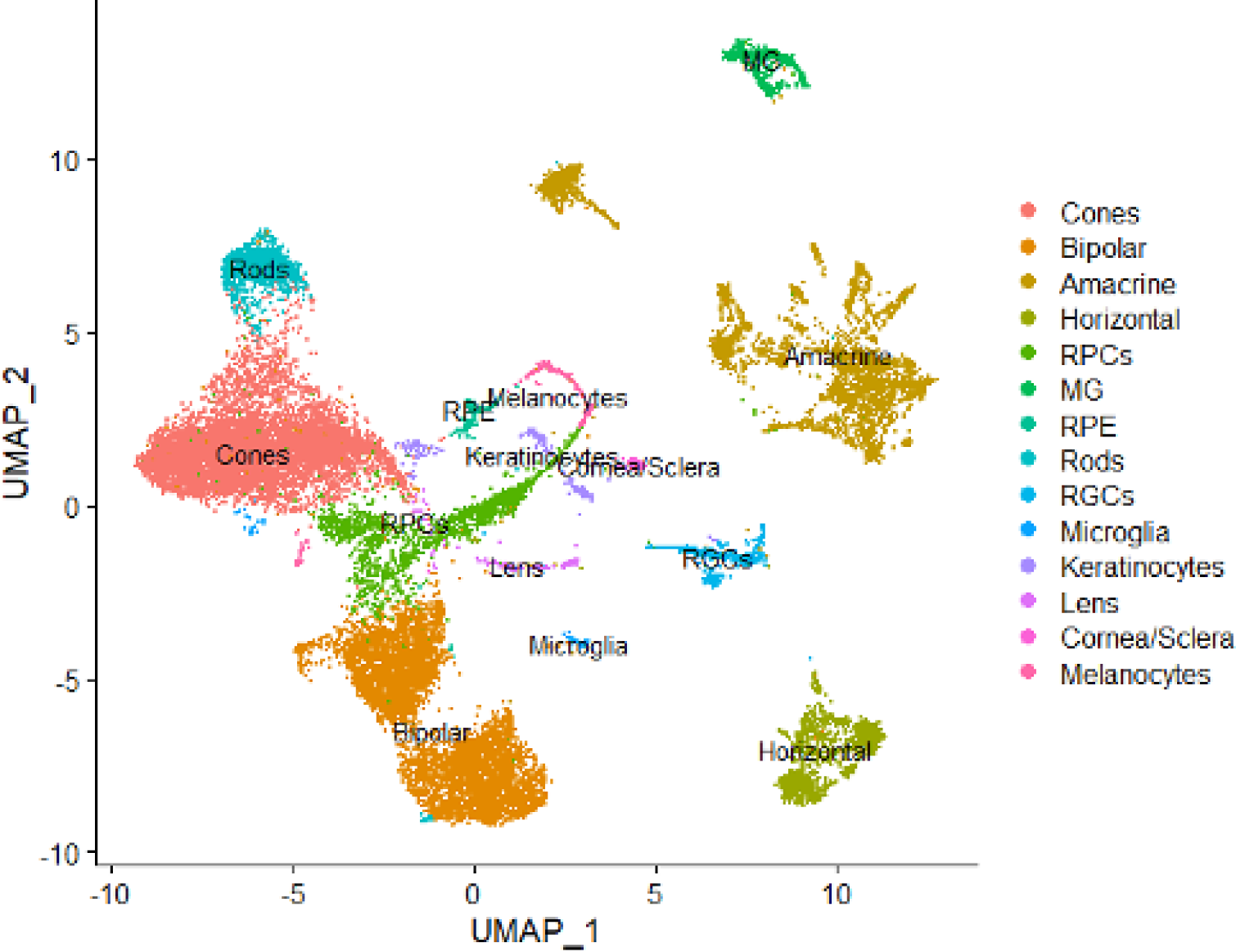
Cell types identified from multiomic sequencing. Identification of cells types in integrated UMAP. Note-each data table is too large to show within word documents so they are uploaded separately in the initial submission, here are all titles and captions for the tables.

**Table S1. DEGs identified in RGC regeneration single-cell RNA sequencing time course**

Differentially expressed genes (DEGs) identified at each timepoint (12, 24, 48, and 72 hpa) between unablated and ablated cells in high clusters of interest (MG, progenitor cells and each retinal neuron type).

**Table S2. Comparison of MG DEGs to widespread injury data**

We compiled lists of all MG DEGs in our data (each timepoint) as well as pooled DEGs from MG data follow widespread LD/NMDA injury from prior studies and identified unique DEGs in each timepoint.

**Table S3. Pseudotime DEGs in RGC regeneration trajectory**

Following isolation of MG, progenitor cells and RGCs, we developed a common pseudotime trajectory from MG>Progenitors>RGCs in unablated and ablated cells. We identified 1,829 DEGs along this trajectory (segmented into 50 timepoints for analysis). Each row contains the gene, the maximum change in expression (delta expression = normalized ablated expression – unablated expression) and the gene ontology cluster each gene belongs to.

**Table S4. Summary of CRISPR/Cas9 screen results**

For each gene tested, we should the average effect size for the gene targeting compared to endogenous RGC regeneration kinetics, adjusted p value compared to control ablated larvae as well as if the gene has a known (K), implicated (I) or unknown role in retinal regeneration from prior work in the field. Genes with a known role are detailed with references(4,8,21,33,35,39,40,68–78).

**Table S5. Pseudotime DEGs in RGC regeneration trajectory in *ascl1a* KO cells**

We generated an analogous pseudotime trajectory with our 24 hpa multiome sequencing data of MG>Progenitors>RGCs. We then identified 269 significant DEGs along the trajectory between *ascl1a* KO and wt ablated only cells.

**Table S6. TFs underlying GRNs in *ascl1a* KO progenitor cells**

Following identification of lineage associated DEGs in progenitor cells at 24hpa between ascl1a KO and wt samples, transcription factors (TFs) that regulate the DEGs were analyzed for differences between samples. Gene regulatory networks (GRNs) were formed based on significantly different TFs between samples.

